# Thermal Control of T-cell Immunotherapy

**DOI:** 10.1101/2020.04.16.045146

**Authors:** Mohamad H. Abedi, Justin Lee, Dan I. Piraner, Mikhail G. Shapiro

## Abstract

Genetically engineered T-cells are being developed to perform a variety of therapeutic functions. However, no robust mechanisms exist to externally control the activity of T-cells at specific locations within the body. Such spatiotemporal control could help mitigate potential off-target toxicity due to incomplete molecular specificity in applications such as T-cell immunotherapy against solid tumors. Temperature is a versatile external control signal that can be delivered to target tissues *in vivo* using techniques such as focused ultrasound and magnetic hyperthermia. Here, we test the ability of heat shock promoters to mediate thermal actuation of genetic circuits in primary human T-cells in the well-tolerated temperature range of 37–42°C, and introduce genetic architectures enabling the tuning of the amplitude and duration of thermal activation. We demonstrate the use of these circuits to control the expression of chimeric antigen receptors and cytokines, and the killing of target tumor cells. This technology provides a critical tool to direct the activity of T-cells after they are deployed inside the body.

## INTRODUCTION

Unlike small molecule and biologic therapies, cells have a natural ability to navigate, persist and proliferate within the body, providing the potential for more targeted and sustained disease treatment. This potential is enhanced by the capacity of cells to probe, process, and respond to their environment and carry out a wide range of sophisticated behaviors, which can be engineered using the tools of synthetic biology^1^. Among the cell types being developed for therapy, T-cells are one of the most promising due to their central roles in cancer, infectious disease and autoimmune disorders, along with their relative ease of isolation, genetic modification and re-engraftment. For example, this potential has been realized in T-cells engineered to express modularly targeted chimeric antigen receptors (CARs), allowing them to specifically eradicate cancers such as lymphomas bearing the CD19 antigen^2,3,4,5^. Unfortunately, it has been challenging to translate these successful results into solid tumors, where CAR T-cells encounter a more immunosuppressive environment^6^ and the risk of sometimes fatal on-target off-tumor toxicity due to the presence of tumor-overexpressed epitopes in healthy tissues^7,8^. Likewise, emerging approaches in which T-cells are used to treat autoimmune disease through local immunosuppression carry the risk of reducing important immune system activity outside the target tissues^9^. Existing strategies seeking to reduce off-target toxicity use additional target recognition elements^10,11^ or chemically triggered kill switches^12,13,14^. However, it can be difficult to ensure perfect recognition solely through molecular markers, and premature termination of T-cell therapy using kill-switches turns off their beneficial therapeutic action.

Here we describe a cellular engineering approach to regulate the activity of therapeutic T-cells with greater specificity through a combination of molecular and physical actuation. This approach is designed to take advantage of the ability of technologies such as focused ultrasound (FUS) and magnetic hyperthermia to non-invasively deposit heat at precise locations in deep tissue^15–18^. By engineering thermal bioswitches allowing T-cells to sense small changes in temperature and use them as inputs for the actuation of genetic circuits, we enable these penetrant forms of energy to spatially control T-cell activity. Our approach is based on heat shock promoters (pHSP), which have been shown to drive gene expression in response to FUS-delivered heating^19–21^, but have not been tested in primary human T-cells. This is important because the behavior of pHSPs varies greatly between cell types and cellular states. In this study, we screen a library of pHSPs in primary T-cells and engineer gene circuits providing transient and sustained activation of gene expression in T-cells in response to brief thermal stimuli within the well-tolerated temperature range of 37–42°C^22–24^. Our circuits incorporate feed-forward amplification, positive feedback and recombinase-based state switches. We demonstrate the use of these circuits to control the secretion of a therapeutic cytokine, expression of a CAR, and killing of target tumor cells.

## RESULTS

### Evaluating candidate pHSPs in primary T-cells

To enable thermal control of T-cell activity, we required a pHSP with robust switching behavior in primary human T-cells. Given the variability in pHSP responses between cell types^25^, we decided to systematically evaluate the activity of 13 different pHSPs in response to a 1-hour incubation at 42°C. This thermal stimulus was chosen based on its tolerability by most tissues^24^, and the convenience of relatively short treatment durations in potential clinical scenarios. Our panel of pHSPs included nine human, three mouse, and one C. *elegans* promoters. The human promoters included four naturally occurring sequences (HSPB, HSPB’2, HSP A/A, HSP A/B), two modifications of HSPB’2 generated by varying the 5’ UTR (HSPB’1, HSPB’3), and three rational modifications of HSPB’2 (SynHSPB’1, SynHSPB’2, SynHSPB’3) inspired by a previously developed sensor of cellular stress^26^. Truncating HSPB’2 and leaving 192 base pairs resulted in SynHSPB’1. To lower potential baseline activity, the AP-1 binding site in SynHSPB’1 was mutated leading to SynHSPB’2. Duplicating SynHSPB’2 four times to increase the number of heat shock elements (HSE) resulted in SynHSPB’3. The three mouse-derived pHSPs were naturally occurring promoters. HSP16, derived from C. *elegans,* was first described in 1986 and is rationally modified to form a minimal bidirectional promoter encompassing four HSE binding sites^27^. HSP16 excludes other transcription factor binding sites that typically exist in human promoters. We incorporated each pHSP into a standardized lentiviral construct in which the pHSP drives the expression of a green fluorescent protein (GFP), with a constitutively expressed blue fluorescent protein (BFP) serving as a marker of transduction (**Fig. 1a**).

**Figure 1.**
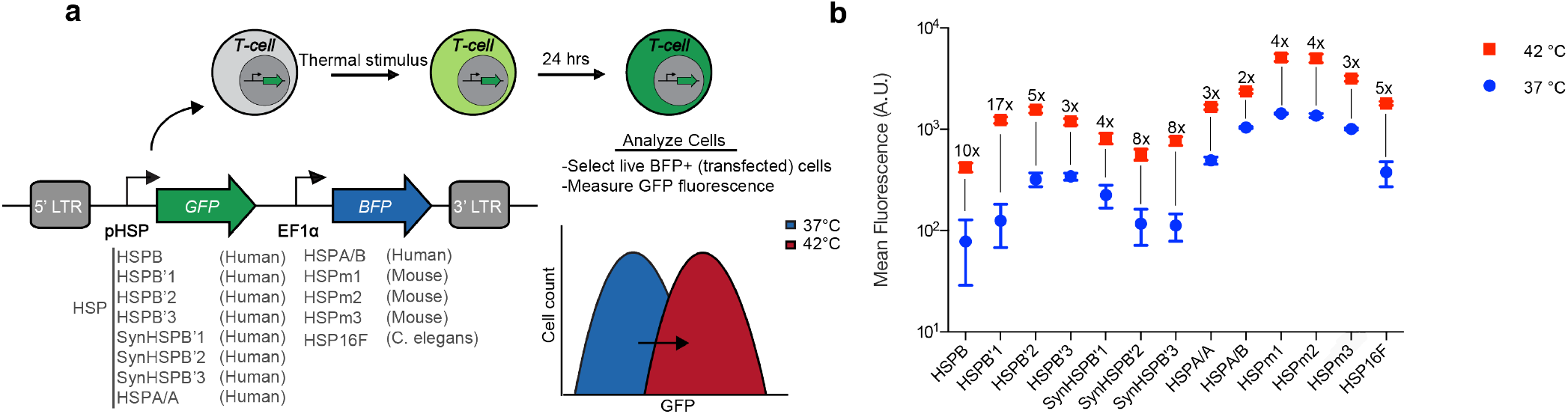
Evaluating candidate pHSPs in primary T-cells. (**a**) Illustration of the screening strategy used to characterize the behaviour of pHSPs. The viral construct used to assay pHSPs is shown, along with the promoters tested. LTR, long terminal repeat. (**b**) Mean fluorescence intensity 24 hours after a 1-hour incubation at 37°C or 42°C, as measured via flow cytometry. The fold change between 37°C and 42°C is listed above each sample. Where not seen, error bars (±SEM) are smaller than the symbol. N=3 biological replicates for each sample.

Of our 13 promoters, HSPB had the lowest baseline expression at 37°C (**Fig. 1b**), an important property for minimizing activity in the absence of the thermal trigger. HSPB’1 showed the largest fold-change in gene expression, reflecting a combination of relatively low baseline expression and strong promoter activity when stimulated. Among the rationally engineered HSPB’2 variants, SYNHSPB’3 had a lower baseline than the natural promoter, albeit with lower maximum expression on activation. The rest of the human and mouse-derived promoters exhibited high baseline activity, resulting in their elimination from further experiments. Finally, the C. *elegans* minimal promoter exhibited acceptable performance and was included in further testing to investigate whether its minimal composition would be advantageous for specific activation in response to temperature. Based on these factors, we chose HSPB, HSPB’1, SynHSPB’3 and HSP16 as our starting points for further circuit engineering.

### Thermal parameters for pHSP activation

After identifying four candidate pHSPs, we tested their response to a range of induction parameters. To search for temperatures that provide rapid induction with minimal thermal burden to the cells, we incubated pHSP-transduced T-cells at temperatures ranging from 37°C to 44°C for 1 hour. All four promoters exhibited a significant increase in activity starting at 42°C (**Fig. 2a**). Increasing the induction temperature beyond this point resulted in a significant enhancement of transcriptional activity, but compromised cell viability (**Fig. 2b**). To optimize induction with minimal cell damage, we chose 42°C for further experiments. We note that unlike the gradual increase in gene expression observed with the mammalian promoters above 42°C, HSP16 exhibited a large jump between this temperature and 43°C, which may make it useful in future circuit engineering applications.

**Figure 2.**
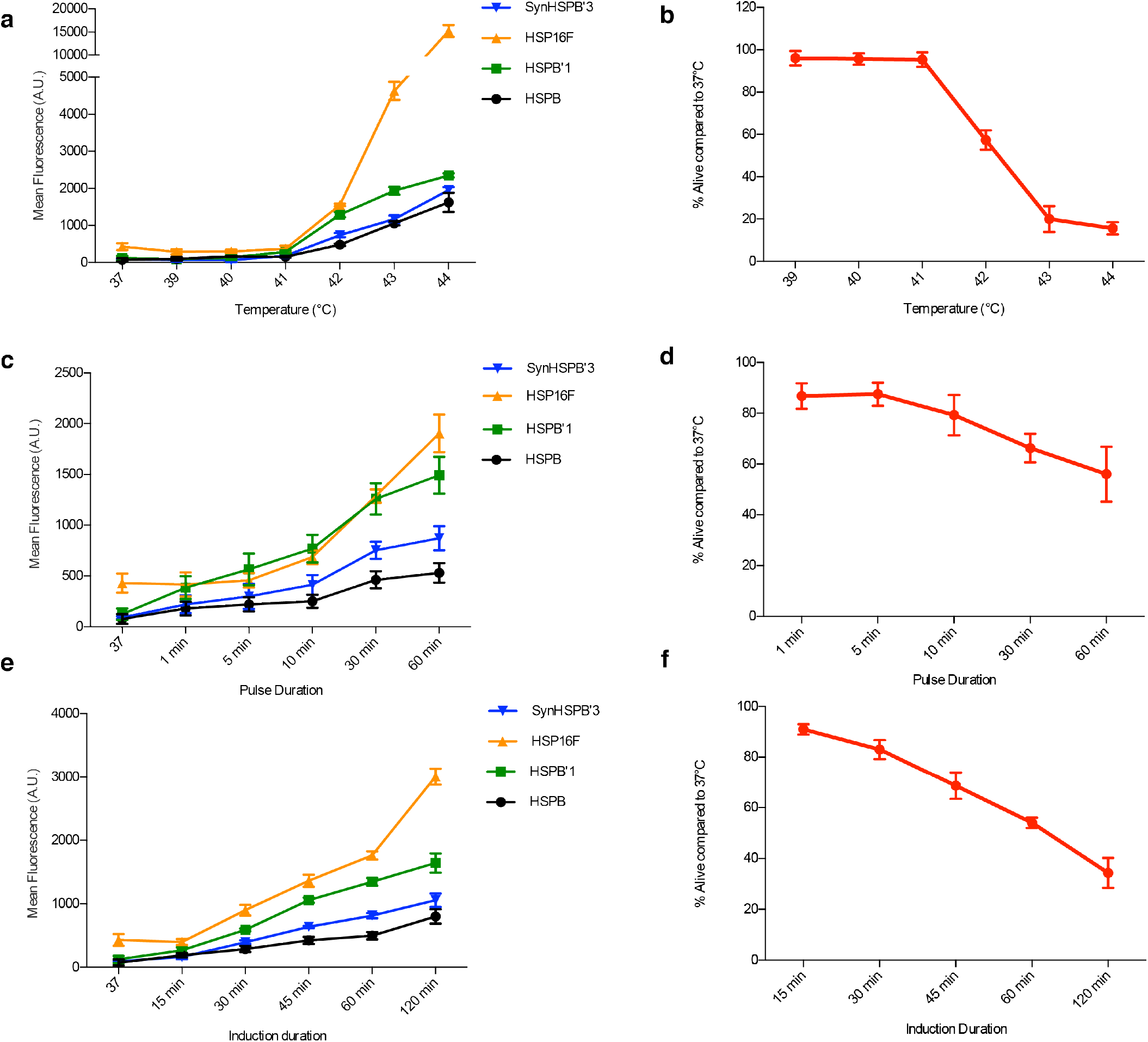
Thermal parameters for pHSP activation. GFP expression from constructs driven by the HSPB, HSPB’1, SynHSPB’3 and HSP16F promoters (**a, c, e**) and T-cell viability (**b, d, f** as a function of **(a,b)** induction temperature for a continuous 1 hour stimulus, **(c,d)** pulse duration of stimuli delivered with a 50% duty cycle alternating between 37°C and 42°C for a fixed thermal exposure of 1 hour, and **(e,f)** induction duration for continuous heating at 42°C. Error bars represent ± SEM. N=3 biological replicates for each sample.

To reduce the effect of thermal exposure on cell viability, we tested a pulsatile heating scheme with a 50% duty cycle^28^. In this scheme, cells underwent repeated cycles of heating to 42°C for a fixed duration and an equal amount of time at 37°C, adding up to a total of one hour at 42°C over a two-hour treatment period. We varied the stimulation period between one minute and continuous heating for 60 min. This experiment revealed a trade-off between promoter activity (**Fig. 2c**) and cell viability (**Fig. 2d**), with shorter pulses reducing the former while increasing the latter. For the purposes of T-cell therapy, in which cells can expand after activation, we decided that a 40% decrease in cell viability was a suitable trade-off for improved activation, therefore selecting a continuous heating paradigm. This paradigm also simplifies the application of heating during therapy. We also investigated continuous stimulation durations ranging from 15 to 120 minutes. Shorter induction enhanced viability (**Fig. 2e**) at the expense of lower gene expression (**Fig. 2f**), with a one-hour stimulation providing the optimal balance. We chose this stimulus paradigm for our subsequent experiments.

### Genetic circuits for amplified and sustained thermal activation

On their own, pHSPs drove a relatively small amount of transient protein expression upon induction. To enable the use of pHSPs in T-cell therapy applications, it is useful to amplify the output of pHSP-driven circuits. This would enable cells to, for example, release a relatively large therapeutic bolus after a single thermal stimulus. To achieve this goal, we implemented a feed-forward amplification circuit in which the pHSP drives an rtTA transactivator, which produces stronger transcriptional activation tunable with doxycycline. In addition, LNGFR was constitutively expressed to identify virally transfected cells (**Fig. 3a**). Amplification circuits incorporating HSPB, HSPB’1, SynHSPB’3 and HSP16 all exhibited a substantial increase in their fold-induction, while only modestly elevating baseline expression. HSPB showed the best performance. To further tune the performance of the HSPB amplifier circuit, we designed constructs with reduced translation of the GFP by varying the Kozak sequence or inserting a micro open reading frame upstream^29^ (**Fig. 3b**). These modifications enabled the tuning of both the baseline expression and the maximal activation level.

**Figure 3.**
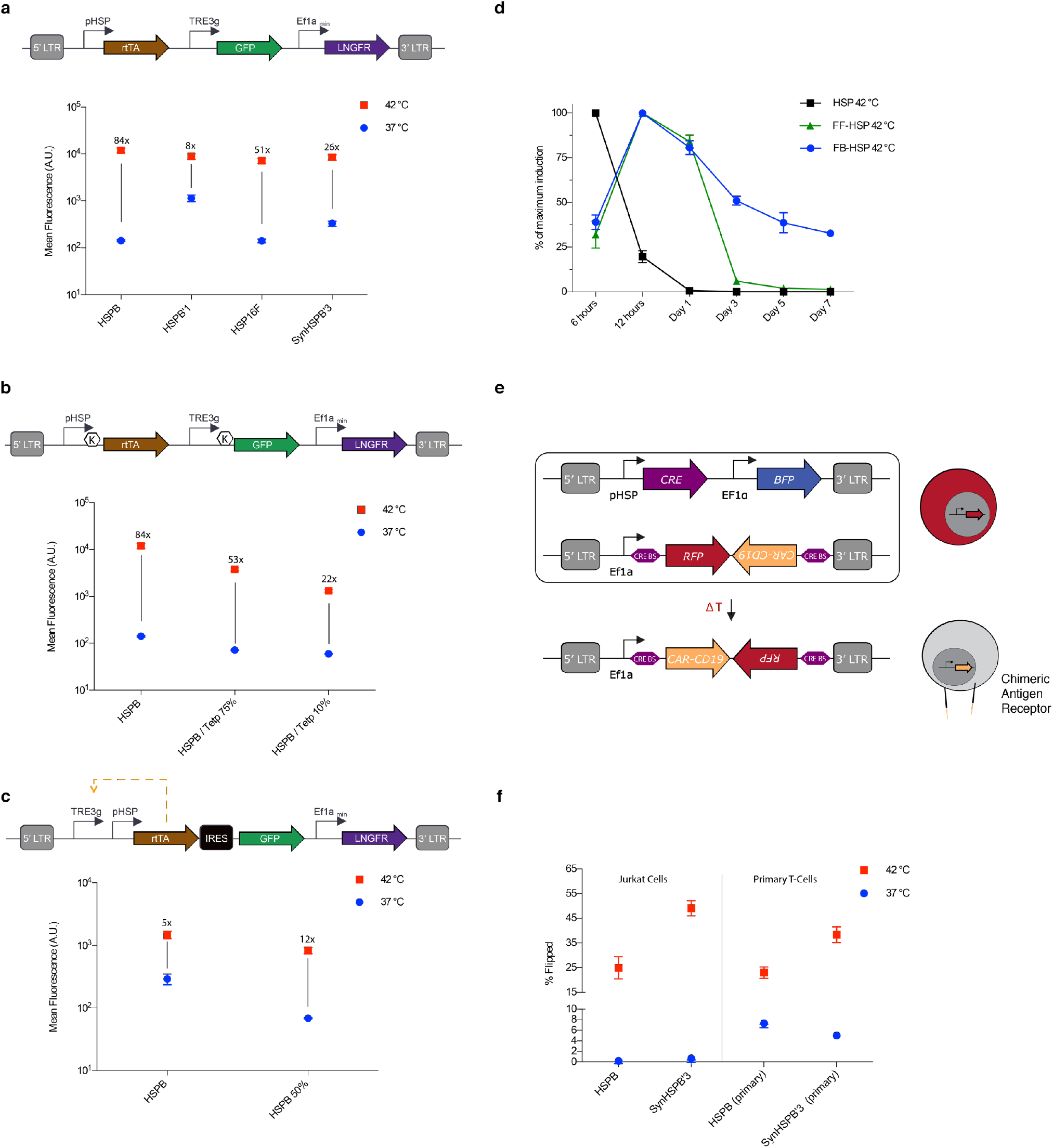
Genetic circuits for amplified and sustained thermal activation. (**a**) Diagram illustrating the thermally trigged feedforward circuit (top). Fluorescence resulting from 1 hour induction at 37° or 42°C for cells supplemented with doxycycline (bottom). (**b**) Diagram illustrating a feed-forward circuit driven by HSPB, <K> indicates varying kozak strength (top). Fluorescence resulting from 1 hour induction at 37° or 42°C for cells supplemented with doxycycline (bottom). (**c**) Diagram illustrating the thermally trigged positive feedback circuit (top). Fluorescence resulting from 1 hour induction at 37° or 42°C for cells supplemented with doxycycline (bottom). (**d**) Normalized expression monitored over seven days after 1 hour induction at 42°C for direct HSPB-driven, feed-forward HSPB and positive-feedback HSPB circuits. Circuits have been modified to replace GFP with a destabilised version of the protein. (**e**) Illustration of the CRE based thermally triggered permanently stable switch designed to express CAR-CD19 upon induction. (**f**) Cells were either incubated at 37°C or thermally stimulated for 1 hour at 42°C and analyzed 24 hours later to determine the number of activated cells. Error bars represent ± SEM. N=3 biological replicates for each sample.

In some therapeutic scenarios, it is critical to prolong the therapeutic action of T-cells following a thermal induction treatment. This would eliminate the need to apply repeated stimuli to maintain treatment efficacy. To develop this capability, we established a positive feedback amplifier circuit by rearranging the elements of our feed-forward amplifier such that rtTA could drive its own expression in the presence of doxycycline (**Fig. 3c**). A similar design was previously tested in human cervical cancer HeLa cells^30^. The HSPB feedback circuit maintained its thermal induction level, and we were able to reduce baseline activity by tuning the Kozak sequence upstream of rtTA. In the current design, the output of the positive feedback circuit is lower than that of the feedforward amplifier, as expected from the GFP payload being placed after an IRES element. While we envision that such “low but steady” activity is desirable in many applications, a “high and steady” mode could in principle be achieved by exchanging the IRES for a 2A element. The dynamic expression profiles of our direct, feed-forward, and feedback HSPB circuits are compared in **Fig. 3d**, demonstrating prolonged expression with positive feedback.

While the positive feedback circuit sustained expression for several days, this circuit can eventually turn off amid dilution or fluctuating expression of the transactivator. To establish a permanent thermal switch, we tested gene circuits in which we placed the expression of CRE recombinase under the control of candidate pHSPs (**Fig. 3e**). In these circuits, the pHSP-driven expression of CRE permanently toggles the circuit from expressing RFP to expressing anti-CD19 CAR by recombining the target vector. When tested in a Jurkat T-cell line, these circuits demonstrated robust activation and minimal leakage (**Fig. 3f**). However, when tested in primary T-cells, we observed significantly higher levels of background activation (**Fig. 3f**). This may arise from the fact that immune stimulation is used to maintain primary T-cells cells in culture and our finding, discussed below, that pHSPs show significant background activity in stimulated primary T-cells. Taken together, these results suggest that in primary T-cells, feed-forward and feed-back amplification provide robust methods for thermal control of gene expression, while pHSP-controlled CRE recombination may produce an unacceptable level of irreversibly accumulating background activation.

### Temperature-activated cytokine release

To demonstrate the ability of our positive feedback circuit to sustain a therapeutically relevant function after thermal induction, we connected its output to the production of a cytokine. The local delivery of cytokines from engineered T-cells would be useful in cancer immunotherapy by allowing T-cells to secrete immune-stimulatory factors to remodel the tumor microenvironment and reduce immunosuppression. It would also be useful in treatments of autoimmune disease by allowing T-cells to secrete factors locally down-regulating the activity of endogenous immune cells. As a model cytokine, we selected IL-21, which has potential utility in cancer immunotherapy due to its ability to stimulate NK cells and CD8^+^ T-cells^31,32^. We incorporated human IL-21 in place of GFP in our positive feedback circuit (**Fig. 4a**). Without thermal induction, primary T-cells transduced with this circuit produced minimal IL-21. Once stimulated, the cells rapidly secreted IL-21, reaching a near-maximal level by 12 hours, and sustained activity for at least 5 days (**Fig. 4a**). The dependence of continued circuit function on doxycycline provides an additional layer of control, allowing the termination of therapy production at a desired time by removing doxycycline. To demonstrate this capability, we removed doxycycline 24 hours after cell induction, resulting in the abrogation of cytokine production by day five. The ability to chemically terminate the activity of our circuit enhances its safety profile in potential therapeutic applications.

**Figure 4.**
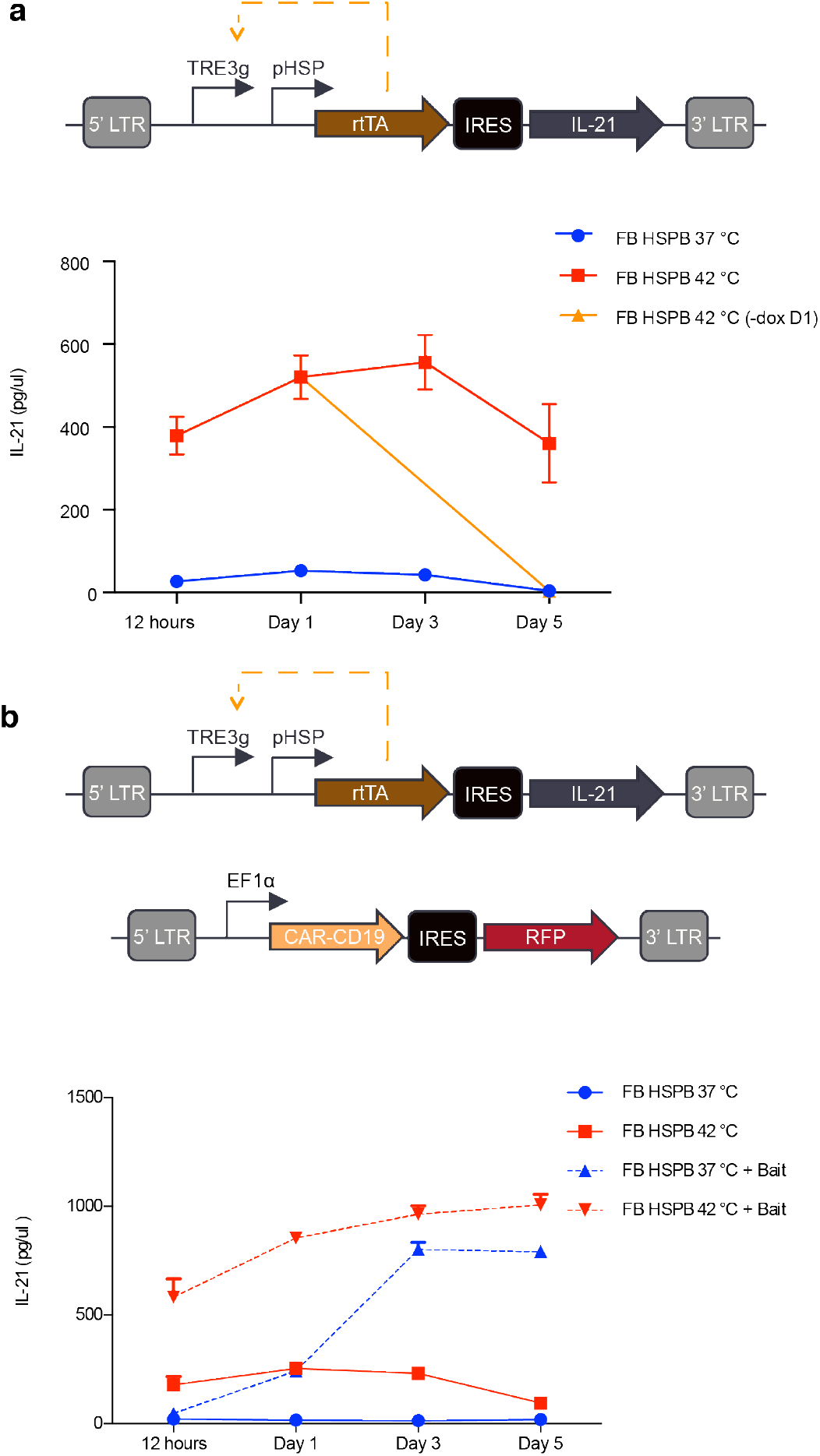
Temperature activated cytokine release. (**a**) Diagram illustrating the positive feedback circuit used to express IL-21 (top). Cumulative IL-21 release from 1-hour induction at 37° or 42°C. In one sample, doxycycline was removed after 24 hours (bottom). (**b**) Illustration of the constructs used to assay the ability of CAR activity to trigger expression of IL-21 in the feedback pHSP circuit (top). Cells were either incubated at 37C or thermally stimulated for 1 hour at 42°C with and without bait cells (bottom). Media was collected and frozen at each time point and all samples were analysed simultaneously at the end of collection. Cumulative IL-21 expression was quantified by using an IL-21 ELISA. Error bars represent ± SEM. N=3 biological replicates for each sample.

In some scenarios, it would be useful for cytokine release to be triggered from a T-cell constitutively expressing a CAR, allowing the cytokine to locally boost immune activation during CAR-directed killing. To test this possibility, we cotransduced primary T-cells with our positive IL-21 circuit and a constitutively expressed anti-CD19 CAR (**Fig. 4b**). In the absence of target Raji bait cells expressing CD 19, IL-21 release was well-controlled by thermal induction (**Fig. 4b**). However, co-incubation with bait cells resulted in the activation of IL-21 release after 3 days in co-culture even in the absence of a thermal treatment (**Fig. 4b**)x. This result is consistent with the induction of pHSP upon T-cell activation, as discussed below.

### Dependence of pHSP-driven circuits on T-cell activation

To directly examine the possibility that pHSPs are turned on in response to CAR-driven T-cell activation, we tested the expression of pHSP-driven GFP in constitutively CAR-expressing T-cells (**Fig. 5a**) upon exposure to a thermal stimulus or bait cells. We found that both thermal stimulation and CAR engagement lead to pHSP-driven gene expression (**Fig. 5b**). This response occurred in cells expressing circuits based on HSPB, SynHSPB’3 and HSPmin promoters. Because SynHSPB’3 lacks the AP-1 site present in wild-type pHSPs such as HSPB, and HSPmin has only HSF1 binding sites, these results suggest that pHSP induction takes place via an HSF1-mediated mechanism. This unexpected finding suggests that activated T-cells experience cellular stress – for example due to rapid proliferation – potentially resulting in an increased number of mis-folded proteins, leading to HSP upregulation. This provides an important insight for the design of thermally inducible immunotherapies.

**Figure 5.**
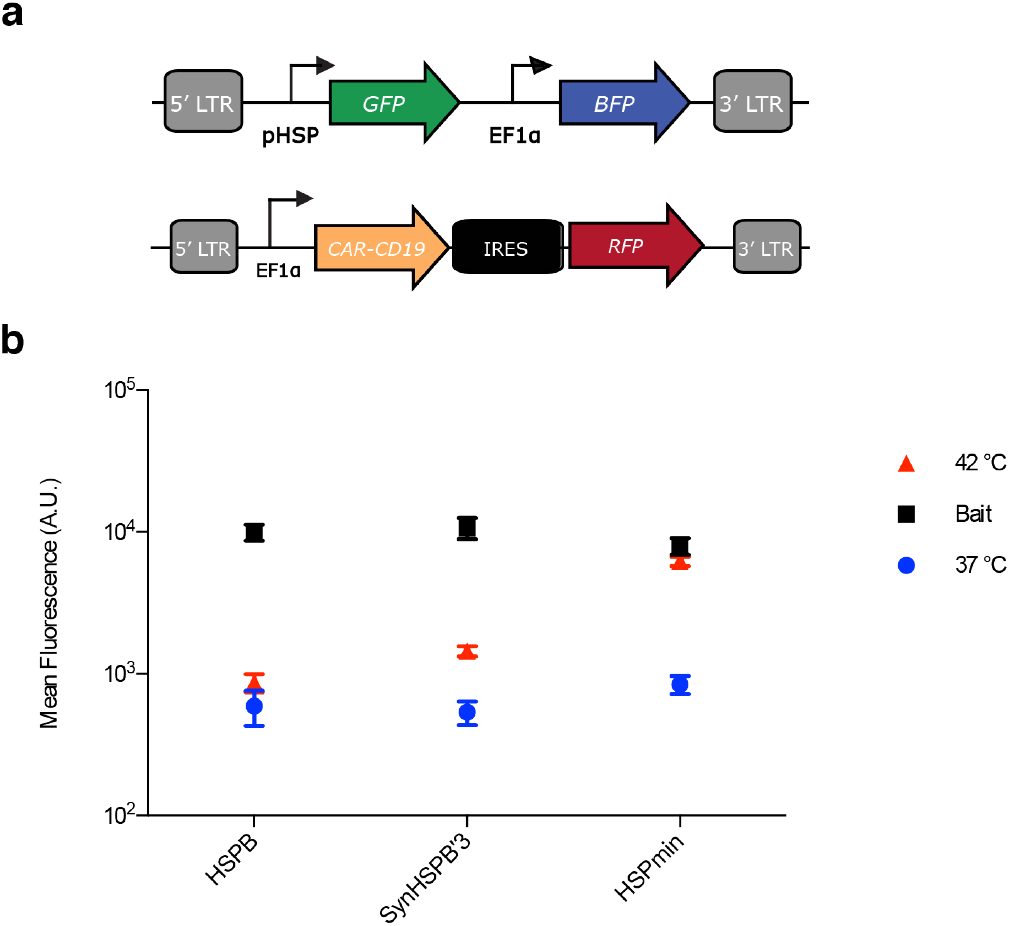
Dependence of pHSP-driven circuits on T-cell activation. (**a**) Illustration of the constructs used to assay the ability of CAR activity to trigger pHSP expression. (**b**) Cells were either incubated at 37°C, thermally stimulated for 1-hour at 42°C, or incubated with CD19^+^ bait cells. pHSP triggered activity was determined by quantifying GFP expression 24-hours after induction. Error bars represent ± SEM. N=3 biological replicates for each sample.

### Auto-sustained thermally induced CAR expression and tumor cell killing

Our finding that CAR engagement drives pHSP activity suggested that a simple, auto-sustained gene circuit could drive CAR-mediated killing in response to the combination of a thermal stimulus and the presence of target cells. In particular, we hypothesized that placing CAR expression under the control of a pHSP (**Fig. 6a**) would result in T-cells with no initial CAR expression or activity, even in the presence of target cells. Upon thermal induction, CAR would become transiently expressed. If the CAR target is present in the vicinity of the T-cells, these cells would become activated, driving sustained expression of additional CAR from the pHSP and target cell killing.

**Figure 6.**
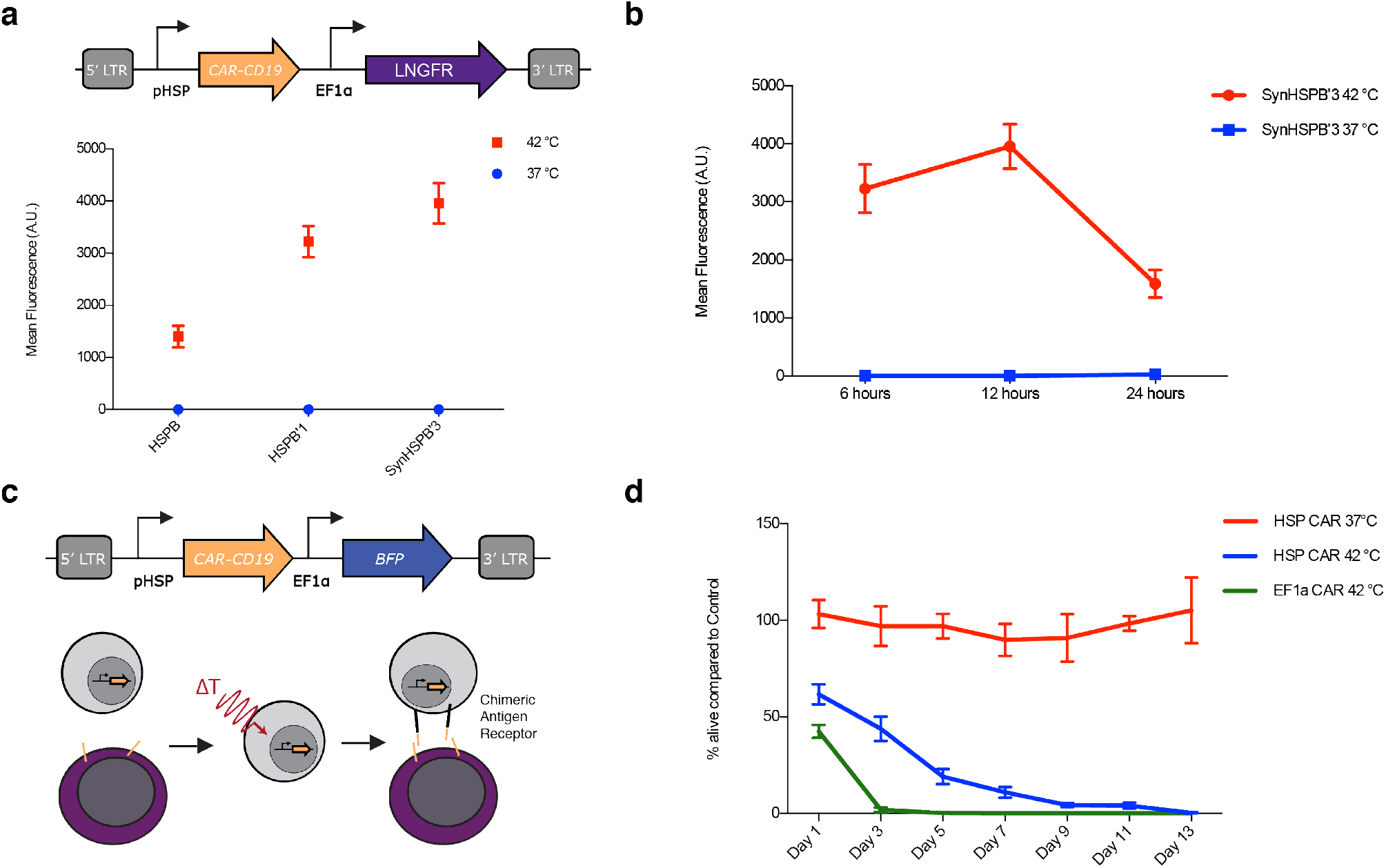
Auto-sustained thermally induced CAR expression and tumor cell killing. (**a**) Illustration of the viral construct used to assay pHSP expression of CAR-CD19. Cells were either incubated at 37°C or thermally stimulated for 1 hour at 42°C and pHSP triggered CAR-CD19 expression was quantified by surface staining of an HA tag appended to the CAR 12 hours after induction. N=3 biological replicates. (**b**) CAR-CD19 expression 6, 12, and 24 hours after 1-hour induction with 37°C or 42°C. N=3 biological replicates. (**c**) Illustration of the viral construct and assay used to test the ability of pHSP inducible CAR expression to conditionally kill bait cells. Cells were either incubated at 37°C or thermally stimulated for 1 hour at 42°C before being incubated with CD19^+^ bait cells. (**d**) Unmodified T-cells and T-cells constitutively expressing CAR-CD19 were used as a negative and positive control respectively. pHSP triggered killing activity was quantified by counting the % of bait cells alive compared to the negative control for a duration of 13 days. N= 3 biological replicates for two T-cell collections from different patients, total N=6. Error bars represent ± SEM.

As predicted, this pHSP-CAR circuit showed no baseline CAR expression in primary T-cells, but began to express CAR when thermally stimulated (**Fig. 6a**). CAR expression was greatly reduced after 24 hours in the absence of target engagement (**Fig. 6b**). When cultured with CD19^+^ bait cells (**Fig. 6c**), thermally activated pHSP-CAR T-cells eliminated the bait cells after 9 days in co-culture (**Fig. 6d**). This killing was as complete as with positive control T-cells carrying a constitutively expressed CAR driven by the EF1α promoter, albeit over a longer time span. When pHSP-CAR T-cells and bait cells were co-incubated without thermal stimulation, no apparent killing took place. These results suggest that a thermal stimulus can kick-start a positive feedback loop of activation-driven expression of CAR from pHSP, leading to effective bait cell elimination.

## DISCUSSION

Our results demonstrate that engineered bioswitch circuits using pHSP can provide control of T-cell therapy with mild hyperthermia. While it has been previously shown that light-switchable proteins could also confer spatiotemporal control over T-cell activity^33^, light has poor penetration into tissues, limiting the utility of such tools. On the other hand, temperature can be elevated at arbitrary depth and with high spatial precision using non-invasive methods such as FUS or magnetic hyperthermia^15–18^.

Our study showed that temperatures in the well-tolerated range of 37-42°C can provide control over T-cell function, including the synthesis and release of a cytokine and the CAR-mediated killing of cancer cells *in vitro*, with minimal baseline activity. In future studies, this performance must be characterized in the *in vivo* setting.

Despite their name, pHSPs can respond to a variety of stimuli such as heat, hypoxia, heavy metals, cytokines and cell division^34,35^. Therefore, the context in which these promoters are being used must be carefully considered. In this work, we capitalized on non-thermal pHSP induction by the T-cell receptor pathway to generate sustained killing circuits. In other contexts where the promiscuous responsiveness of pHSPs presents an un-exploitable hindrance, it may be desirable to develop thermal response mechanisms based on orthogonal molecular bioswitches^22,23^.

## MATERIALS AND METHODS

### Plasmid Construction and Molecular Biology

All plasmids were designed using SnapGene (GSL Biotech) and assembled via KLD mutagenesis or Gibson Assembly using enzymes from New England Biolabs. After assembly, constructs were transformed into NEB Turbo and NEB Stable E. coli (New England Biolabs) for growth and plasmid preparation. The CAR-CD19 gene containing the CD28 and CD3z signaling domain was a kind gift from the Laboratory of David Baltimore (Caltech). Integrated DNA Technologies synthesized other genes, the pHSP, and all PCR primers.

### Cell Lines

Raji cells (CCL-86) were obtained from ATCC and cultured in RPMI 1640 media (Thermo Fisher Scientific) with 1x Penicillin/Streptomycin (Corning). GFP^+^ Raji cells were constructed via viral infection of a GFP driven by the EF1a promoter. Lentivirus was prepared using a third-generation viral vector and helper plasmids (gifts of D. Baltimore). Virus was packaged in HEK293T cells grown in 10 cm dishes. After 3 days of transfection, viral particles were concentrated via Ultracentrifugation. Infection was performed by following the “RetroNectin” reagent protocol. Experiments were performed at least two weeks after infection.

### Primary T-cells

T-cells were isolated with the EasySep Human T-cell isolation Kit (STEMCELL Technologies) from frozen human peripheral blood mononuclear cells obtained from healthy donors. T-cells were stimulated with CD3/CD28 Dynabeads (Thermo Fisher Scientific) at 1:1 cell:bead ratio for 1 day before viral transduction. T-cells were cultured in RPMI supplemented with 50 U/ml IL-2 (Miltenyi Biotech) and 1 ng/ml IL-15 (Miltenyi Biotech) every other day. Dynabeads were removed after 7 days of culture. T-cells were enriched by LNGFR magnetic bead based sorting (Miltenyi Biotech) when appropriate.

### Thermal Regulation Assay

Thermal stimulation of T-cells was performed in a Bio-Rad C1000 thermocycler. T-cells at 1-2 million/ml were supplemented with doxycycline, if needed, and mixed well before transferring 50 μl into a sterile PCR tube. The temperature and duration of stimulation was varied based on the experimental procedure. Upon completion of thermal stimulation, cells were moved back into a mammalian incubator and supplemented 1:1 with fresh media containing cytokines and in some cases doxycycline. Cells were typically incubated for 24 hours unless stated otherwise before assaying with a flow cytometer (MACSQuant VYB). Dead cells were typically excluded via FSC/SSC gating for routine assays. In figure 2, a LIVE/DEAD viability/cytotoxicity kit (Thermo Fisher) was used for a more accurate quantification of cell state. Live cells were further gated via a fluorescent protein or LNGFR stain to isolate virally infected cells for further analysis. The change in mean fluorescence of the cell population was used to characterize the fold change of pHSP constructs. Anti-HA antibodies (Miltenyi Biotech) were used to stain for CAR expression and V450 Mouse Anti-human CD271 was used to stain LNGFR. IL-21 expression was measured using a human IL-21 DuoSet ELISA (R&D systems).

### T-cell Bait Assay

Raji and GFP^+^ Raji cells were used as bait cells for CAR-CD19 T-cells. Bait assays were initiated by mixing T-cells with bait cells at a 3:1 ratio. This ratio was established to avoid excessive bait cell growth before T-cell engagement. To assess T-cell killing of bait cells GFP^+^ Raji were used and the count of GFP^+^ cells was tracked over time.

### Data and code availability

Plasmids will be made available through Addgene upon publication. All other materials and data are available from the corresponding author upon reasonable request.

## ACKNOWLEDGEMENTS

The authors thank Ellen Rothenberg, David Baltimore, Arnab Mukherjee, and Yvonne Chen for helpful discussions. M.H.A. was supported by the NSF graduate research fellowship and the Paul and Daisy Soros Fellowship for New Americans. This research was funded by the Sontag Foundation and the DARPA Young Faculty Award. Related research in the Shapiro laboratory is supported by the Burroughs Wellcome Career Award at the Scientific Interface, the Packard Fellowship in Science and Engineering and the Heritage Medical Research Institute.

## AUTHOR CONTRIBUTIONS

M.H.A. and M.G.S. conceived the study. M.H.A., J.L. and D.I.P. planned and performed experiments. M.H.A. and J.L. analyzed data. M.H.A. and M.G.S. wrote the manuscript with input from all other authors. M.G.S. supervised the research.

### Competing interests

The authors declare no competing financial interests.

## Notes

### Competing Interest Statement

The authors have declared no competing interest.

